# Enhancing Recombinant Vector Assembly Efficiency: A Novel Methodological Approach

**DOI:** 10.64898/2026.07.30.741931

**Authors:** Shu Yuan, Hailong Jiang, Haiyan Wang, Mengmeng Fu, Juan Wang, Zhen Liu, Yang Li

**Affiliations:** Biorun Research, Wuhan BioRun Biosciences Co., Ltd., Wuhan, Hubei 430070, China

**Keywords:** Recombinant vector, Construction efficiency, DNA processing, Gene cloning, Gene therapy

## Abstract

With the rapid development of modern biotechnology, DNA vectors have become fundamental tools for inserting, transferring, and expressing specific gene sequences in various fields such as gene cloning, gene expression, gene editing, and gene therapy. However, when dealing with complex structured DNA sequences, traditional vector construction methods face challenges with low connection efficiency. This study proposes a new method for constructing recombinant vectors by employing a strategy of high-temperature treatment followed immediately by placement on ice, effectively reducing the complexity of DNA structures and enhancing the efficiency of PCR product-vector connection, thereby improving the construction efficiency of recombinant vectors. This paper describes the technical details of the method, experimental validation, and applications in gene cloning, gene recombination editing, and the preparation of gene therapy drugs, providing a new efficient tool for molecular biology experiments.

## Introduction

DNA vectors, commonly referred to as plasmids or recombinant vectors, are pivotal tools in biotechnology for the insertion, transfer, and expression of specific gene sequences. They are indispensable in gene cloning, gene expression, gene editing, and gene therapy, where they serve as carriers for foreign DNA into host cells. With advancements in gene research and regulatory mechanism studies, scientists increasingly face challenges when dealing with complex DNA sequences, such as those with repetitive elements, high GC content, long fragments, or other structural complexities. These sequences often present obstacles during DNA replication, transformation, and subsequent applications like PCR amplification, cloning, and vector insertion (Sambrook & Russell, 2001). The structural integrity and stability of such sequences can negatively affect the efficiency of gene cloning, thus reducing the success rates of molecular biology experiments (Mullis et al., 1986).

Traditional methods of recombinant vector construction, including enzymatic ligation and homologous recombination, have long been used to assemble DNA fragments with vectors. However, these techniques are often inefficient when dealing with complex DNA structures, sometimes resulting in low connection efficiency or even failure to join the DNA fragments to vectors (Sambrook & Russell, 2001; Dower et al., 1988). The challenges associated with these methods underscore the need for optimized approaches that can address the unique issues posed by complex DNA sequences. Such methods would increase the likelihood of successful DNA-vector ligation, thereby enhancing the efficiency of downstream experiments and improving overall success rates in gene manipulation and cloning (Green & Sambrook, 2012). Recombinant vectors have evolved into essential tools in molecular biology, enabling precise gene manipulation for both fundamental research and applied biotechnology. The concept of recombinant vectors was first significantly advanced in the early 1980s, with the pioneering work of Hanahan (1983), who developed plasmid vectors for the transformation of *E. coli*, laying the foundation for gene expression studies. Subsequent developments in the 1980s and 1990s expanded the utility of these vectors, exploring various vector systems for gene cloning in *E. coli* and other organisms. Jasny et al. (1984) investigated the use of plasmids and bacteriophages in gene cloning, while Vojta and Tullius (1987) provided early insights into vector construction techniques. These early innovations formed the basis for modern recombinant DNA technologies.

The progress of recombinant vector technology has led to its widespread application in synthetic biology, with notable breakthroughs such as the work by Gibson et al. (2010), who demonstrated the assembly of synthetic bacterial genomes using recombinant vectors. This method highlighted the potential of vectors for constructing complex genetic constructs and advancing synthetic biology. Similarly, Luo et al. (2002) introduced new expression vectors designed to optimize protein yield in *E. coli*, while Zhang et al. (2011) developed versatile plasmid systems that enable efficient protein expression across various bacterial species. These advancements have facilitated the production of recombinant proteins for research and industrial applications.

Moreover, recombinant vectors have expanded beyond prokaryotic systems, demonstrating their versatility in mammalian cells. Rosamond and Hileman (1989) explored the use of plasmid-based vectors for gene expression in both bacterial and mammalian systems, underscoring the broad applicability of recombinant vectors in genetic engineering. Collectively, these innovations have significantly enhanced the ability to clone, express, and manipulate genes, not only advancing our understanding of genetics but also providing practical applications in medicine, agriculture, and biotechnology (Cohen et al., 1973; Morrow et al., 1983).

Therefore, continuously optimizing the methods for constructing recombinant vectors is vital for enhancing the efficiency and success rate of gene cloning and related experiments, particularly when dealing with complex DNA structures. By improving the connection between DNA fragments and vectors, these advancements will provide researchers with more reliable tools for studying and engineering genes, thus enabling progress in a wide range of scientific disciplines.

## Methods

### Separate various plasmid types based on their mobility rates

1.0 g of agarose (Wuhan BioRun Biosciences Co., Ltd.) was weighed and placed into a conical flask, followed by the addition of 100 ml of 1x TAE electrophoresis buffer. The mixture was heated in a microwave until the agarose was completely dissolved. After cooling to approximately 50–60°C, the solution was gently mixed and poured into a gel casting mold with a comb inserted. The gel was allowed to solidify. An appropriate volume of pGADT7 plasmid (8393 bp) solution was mixed with an adequate amount of 6x loading buffer to ensure thorough mixing. The solidified gel was placed into the electrophoresis tank and submerged with 1x TAE buffer until the liquid level covered the gel. The mixed samples were carefully loaded into the wells of the gel, and DNA Marker 5000 (Wuhan BioRun Biosciences Co., Ltd.) was added to the adjacent wells as a molecular weight standard. The power supply of the electrophoresis apparatus was connected, and the voltage was set at 180 V for approximately 12 min. After electrophoresis, the gel was transferred to a gel imaging system for observation and photography under UV light. The size and conformation of the plasmid were determined based on the band positions of the DNA Marker.

### Construction of pSYB02 Vector (DNA ligase method)

A 2652 bp fragment was amplified using the plasmid pGreenII 0800-LUC-ccdb as a template with primers OSYB1 (CCGGATCCCCACTAGCCTTGACAGGATATAT) and OSYB2 (CCAAGCTTCTTTGGTCTTCTGAGACTGT) in a 2x 50 μL reaction system. The target fragment was then purified by gel extraction. Heat Treatment Group: The PCR product was subjected to heat treatment at 95℃ for 3 min, followed by immediate cooling on ice for 2 min. Control Group: The PCR product was not subjected to heat treatment. The PCR products were digested with *Hind* III and *BamH* I at 37℃ for 2h. The digested products were run on a 1% agarose gel and photographed in high-definition TIFF format. The target bands were excised from the gel and purified using a gel extraction kit.

Primers OSYB3 (AGCTTAGGGCCCTCCTAGGCACTAGTATCTAGAAGGTACCAGAGCTCCG) and OSYB4 (GATCCGGAGCTCTGGTACCTTCTAGATACTAGTGCCTAGGAGGGCCCTA) were annealed using the following conditions: 95℃ for 3 min, followed by a temperature decrease to 22℃ over 10 min. The purified fragment was ligated to the annealed product using T4 DNA Ligase in a 10 μL reaction system.

5 μL of the ligation mixture was transformed into 50 μL of competent DH5α cells. The competent cells were retrieved from the -80℃ freezer and thawed on ice. The recombinant product was added to the competent cells and gently mixed. The mixture was placed on ice for 30 min, followed by heat shock at 42℃ for 90 sec. Immediately after heat shock, the tube was placed on ice to cool for 2 min. The cells were then resuspended in 500 μL of antibiotic-free LB liquid medium and incubated at 37℃ with shaking at 150-200 rpm for 60 min. The transformed cells were plated on LB solid medium containing kanamycin and incubated at 37℃ for colony formation. The following day, the number of positive monocolonies were compared between the heat-treated group and the control group.

### Construction of pSYB-Dual-LUC01-TBF1 (General fragment)

TBF1-1: The 483 bp TBF1-5’UTR fragment was amplified using primers OSYB24 (CCGGTCTCCGATATTCTAGAAACAGCATCCGTTTTTA) and OSYB25 (CCGGTCTCCCCCGTCTCCGGCGAACTTTTTTTATTTTA) with Arabidopsis<colcnt=3> genomic DNA (gDNA) as the template. This fragment was designed for subsequent ligation (enzyme digestion and ligation) .TBF1-2: The 483 bp TBF1-5’UTR fragment was amplified using primers OSYB26 (CTCGAGGTCGACGGTATCTCTAGAAACAGCATCCGTTTTTA) and OSYB27 (GGCGTCTTCCATGGTCCCTCCGGCGAACTTTTTTTATTTTA) with Arabidopsis gDNA as the template. This fragment was designed for homologous recombination. The parent vector pSYB-Dual-LUC01 was retrieved from the -80℃ freezer and subjected to *Bsa*I digestion. The TBF1-1 fragment was also processed with *Bsa*I digestion.

Control Group 1: Followed the above procedures without additional treatments. Control Group 2: The TBF1-1 fragment, TBF1-2 fragment, and pSYB-Dual-LUC01 were subjected to heat treatment at 95℃ for 3 min and then allowed to cool naturally. Experimental Group: The TBF1-1 fragment, TBF1-2 fragment, and pSYB-Dual-LUC01 were subjected to heat treatment at 95℃ for 3 min, followed by immediate cooling on ice for 2 min. The *Bsa*I-digested TBF1-1 fragment was ligated to the *Bsa*I-digested pSYB-Dual-LUC01 vector using T4 DNA Ligase. The ligation mixture was then transformed into competent Top10 cells using electroporation.

The TBF1-2 fragment was recombined with the *Bsa*I-digested pSYB-Dual-LUC01 vector using a homologous recombination kit. The recombination mixture was then transformed into competent Top10 cells using electroporation. The transformed cells were plated on LB solid medium containing kanamycin and incubated at 37℃ for colony formation.

### Construction of pSYB06 Vector (Complex fragment)

A 396 bp fragment of the EIL1-5’UTR was amplified using Arabidopsis cDNA as the template. Fragment EIL1-1 was amplified with OSYB20 (CCGGTCTCCGATATTGCTCATTTATTTATTATATATTTTGG) and OSYB21 (CCGGTCTCCCCCGTGTCTCTTCCACCACAATCAAG) for subsequent ligation. Fragment EIL1-2 was amplified with OSYB22 (CTCGAGGTCGACGGTATCTGCTCATTTATTTATTATATATTTTGG) and OSYB23 (GGCGTCTTCCATGGTCCGTCTCTTCCACCACAATCAAG) for homologous recombination. The parent vector pSYB-Dual-LUC01 was retrieved from the -80℃ freezer and subjected to *Bsa*I digestion. The EIL1-1 fragment was also processed with *Bsa*I digestion.

Treatment Group: The EIL1-1 fragment, EIL1-2 fragment, and pSYB-Dual-LUC01 were subjected to heat treatment at 95℃ for 3 min, followed by immediate cooling on ice for 2 min. Control Group 1: Followed the above procedures without heat treatment. Control Group 2: The EIL1-1 fragment, EIL1-2 fragment, and pSYB-Dual-LUC01 were subjected to heat treatment at 95℃ for 3 min and then allowed to cool naturally. The *Bsa*I-digested EIL1-1 fragment was ligated to the *Bsa*I-digested pSYB-Dual-LUC01 vector using T4 DNA Ligase. The ligation mixture was then transformed into competent Top10 cells using electroporation. The EIL1-2 fragment was recombined with the *Bsa*I-digested pSYB-Dual-LUC01 vector using a homologous recombination kit. The recombination mixture was then transformed into competent Top10 cells using electroporation.

### RNA structure prediction

EIL1 fragment (396 bp) was predicted with the complex secondary structure using RNAfold (http://rna.tbi.univie.ac.at/cgi-bin/RNAWebSuite/RNAfold.cgi).

## Results

The influence of plasmid type and structure on migration rates during agarose gel electrophoresis is illustrated in the data, showing how different plasmid conformations affect their movement through the gel (Fig 1). Fig 1a shows the experimental data, demonstrating that plasmids with a supercoiled structure exhibit the highest mobility, followed by relaxed circular plasmids, while linear plasmids display the slowest migration. These results suggest that plasmid conformation—supercoiled, relaxed, or linear—affects the rate at which they move through the gel, with supercoiled plasmids being the most compact and thus migrating more efficiently. Fig 1a depicts a schematic representation of the three plasmid types, highlighting the structural differences among supercoiled, relaxed, and linear forms. This variability in plasmid structure may have important implications for downstream applications such as cloning, transfection, or expression studies. Furthermore, these results suggest that similar structural variations could exist in other forms of double-stranded DNA, which may also influence their behavior in experimental settings, particularly in terms of mobility and interaction with other molecules.

**Figure 1.**
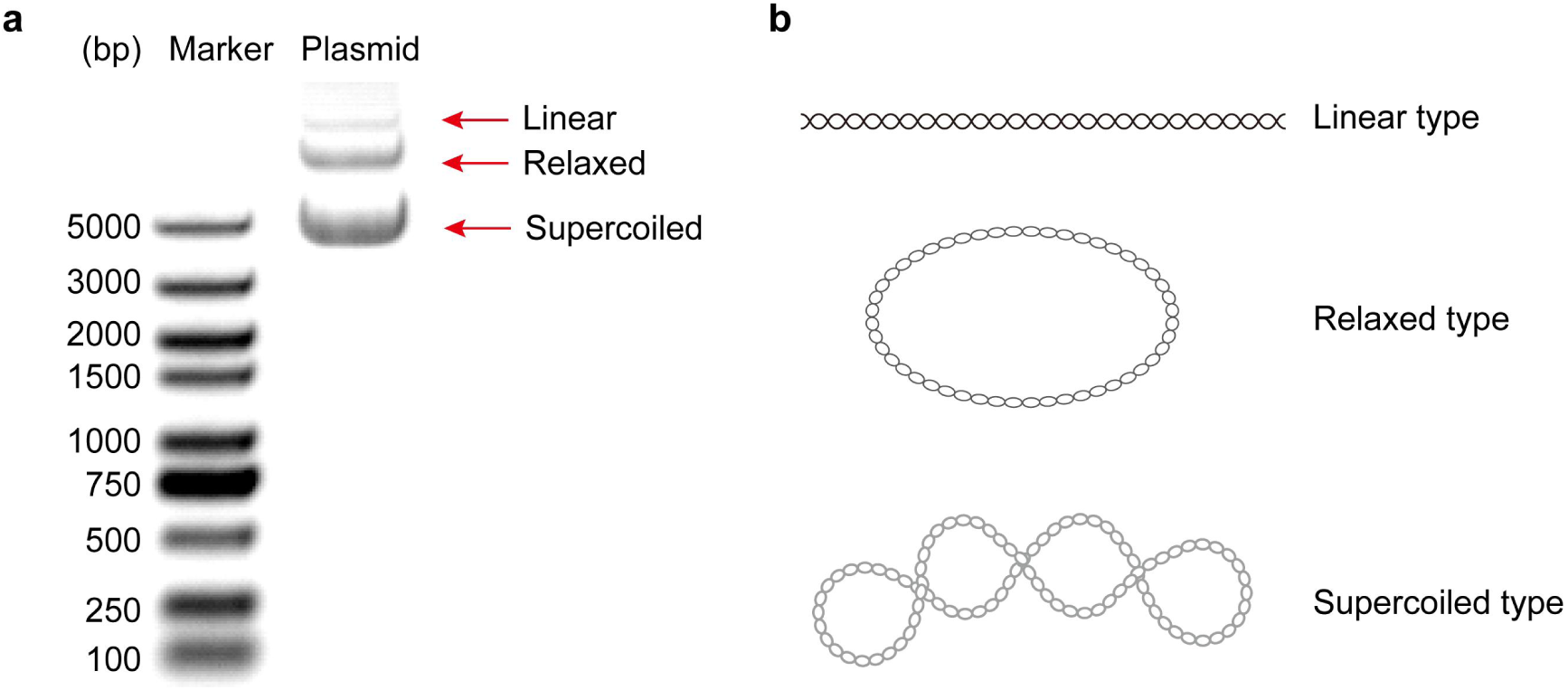
Influence of plasmid type and structure on the mobility rate. This figure illustrates the effect of different plasmid types and their structural configurations on the rate of movement through agarose gel electrophoresis. (a) Experimental data showing the migration rate of pGADT7 plasmids through agarose gel electrophoresis. Data demonstrate that plasmids with a supercoiled structure exhibit the highest mobility, followed by relaxed circular forms, while linear plasmids display the slowest migration. (b) Schematic representation of plasmids with varying structural characteristics, including linear, relaxed, and supercoiled types.

The effect of heat treatment on the digestion efficiency of a 2652 bp PCR product with *Hind*III and *BamH*I restriction enzymes was evaluated (Fig2). Agarose gel electrophoresis (Fig 2a) revealed that heat treatment led to a more defined digestion pattern, with discrete bands indicating improved cleavage efficiency. In contrast, the untreated sample showed smeared bands, suggesting incomplete digestion. These observations point to the enhanced specificity and efficiency of the enzyme digestion following heat treatment. Additionally, the impact of heat treatment on monoclonal colony formation (Fig 2b) was assessed, with heat-treated samples yielding a higher frequency of colonies compared to untreated controls. This further supports the conclusion that heat treatment promotes more efficient and specific enzyme digestion, which may improve the success of downstream cloning and molecular experiments.

**Figure 2.**
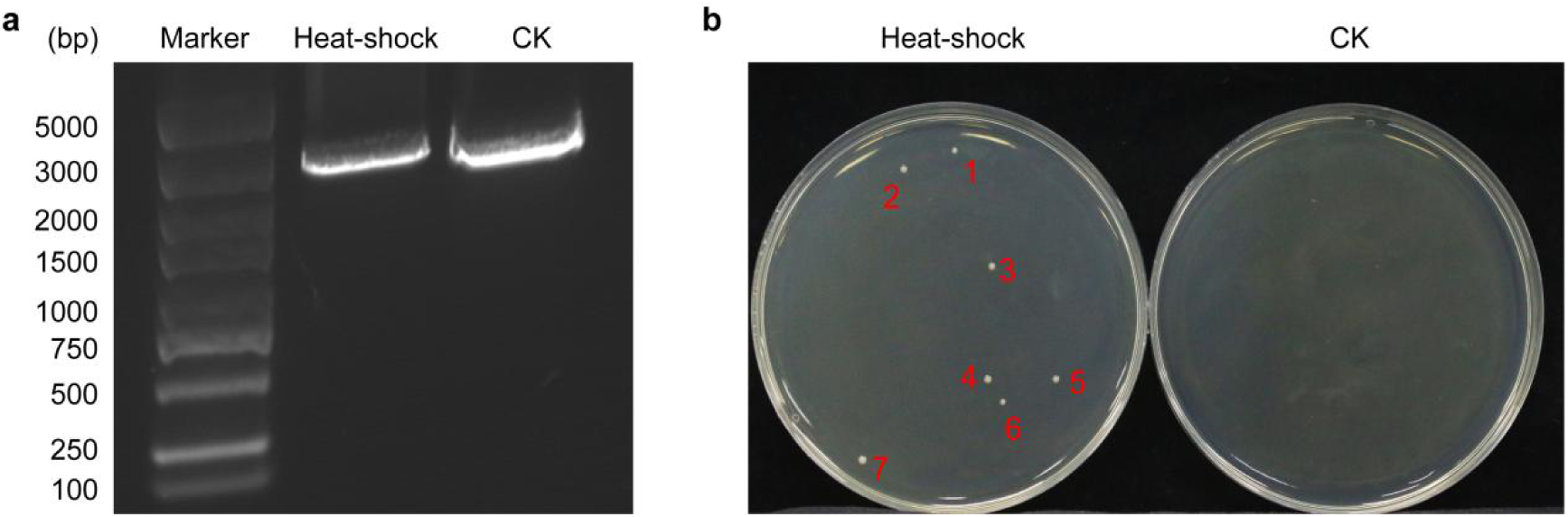
Effect of heat treatment on the digestion of PCR products with restriction enzyme enzymes. This figure demonstrates the impact of heat treatment on the digestion efficiency of a 2652 bp PCR product using *Hind*III and *BamH*I enzymes. (a) Agarose gel electrophoresis showing the digestion pattern of the PCR product with *Hind*III and *BamH*I, with or without heat treatment. The heat-treated digestion results in more discrete bands compared to the untreated, indicating improved digestion efficiency. (B) Comparison of monoclonal colony formation following digestion with and without heat treatment. The heat-treated samples show a higher frequency of colonies (pSYB02 vector), suggesting more efficient and specific cleavage of the PCR product.

The effect of heat-shock treatment on monoclonal colony formation for recombinant vectors was assessed under three different conditions: CK1 (non-treated), CK2 (heat-shock only, without chilling), and a full heat-shock treatment (including chilling) (Fig 3). The experiment compared vectors constructed via DNA ligase and homologous recombinase methods. The results demonstrate that vectors subjected to the full heat-shock treatment consistently yielded a higher number of monoclonal colonies than those in the CK1 and CK2 groups, regardless of the construction method used. This suggests that the inclusion of a chilling step, in addition to heat-shock, significantly improves the efficiency of monoclonal colony formation. These findings indicate that heat-shock treatment, particularly with chilling, enhances recombinational efficiency, making it a critical step for improving the success of recombinant vector applications.

**Figure 3.**
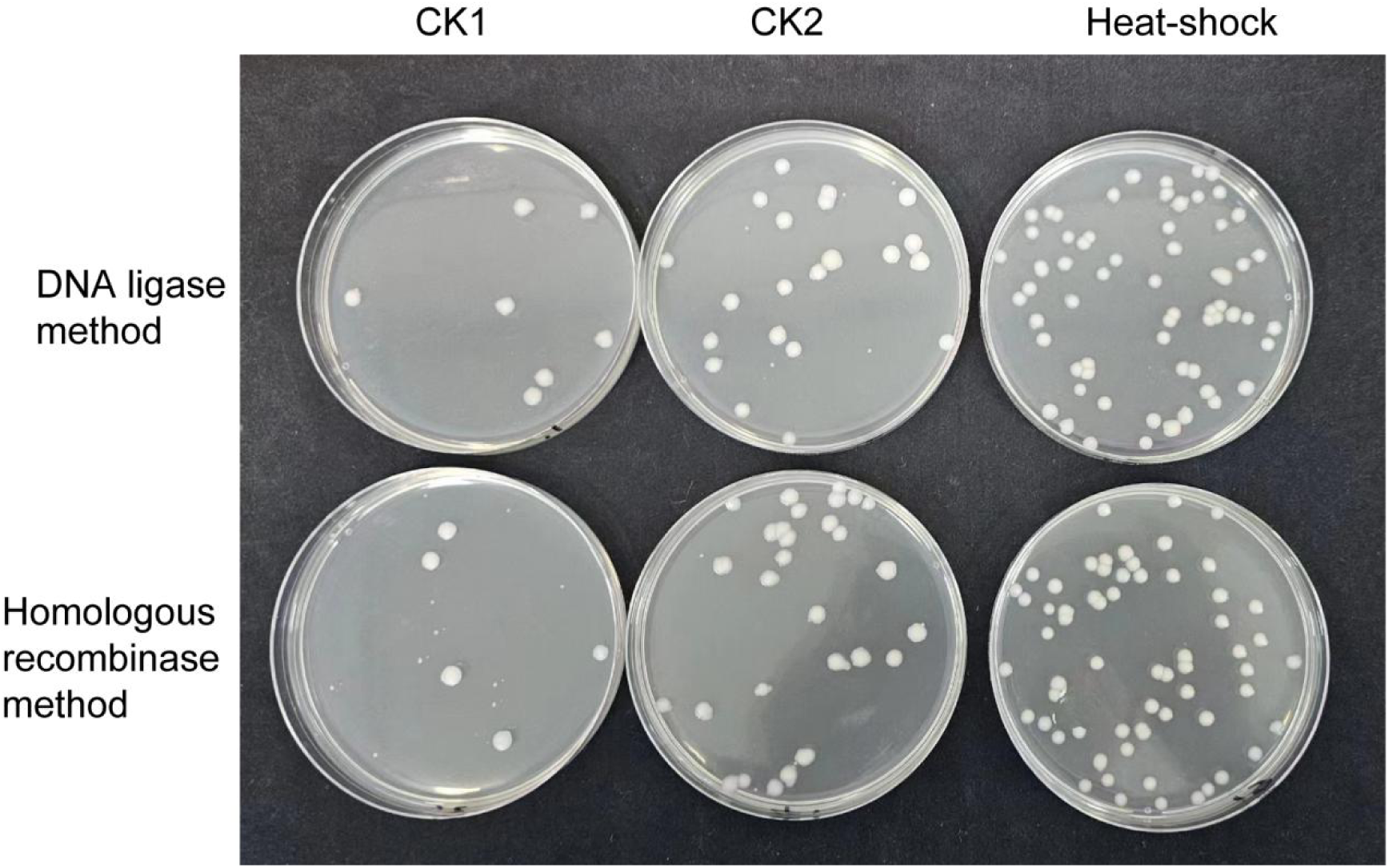
Effect of heat-shock treatment on monoclonal colony formation for recombinant vectors (pSYB-Dual-LUC01-TBF1). This figure illustrates the impact of different treatments on the efficiency of monoclonal colony formation for general recombinant vectors. The experiments compared vectors constructed via DNA ligase or homologous recombinase methods. Three conditions were tested: CK1 (non-treated), CK2 (heat-shock only, without chilling), and heat-shock treatment (including chilling). The results show that vectors subjected to heat-shock treatment (including a chilling step) consistently yield more monoclonal colonies than CK1 and CK2, regardless of the vector construction method. Data indicate that the heat-shock treatment significantly enhances recombinational efficiency.

The impact of heat-shock treatment on monoclonal colony formation for complex recombinant vectors was evaluated using both DNA ligase and homologous recombinase construction methods (Fig 4). Fig 4a presents the RNAfold-predicted secondary structure of the *EIL1* fragment (From arabidopsis), which reveals a highly intricate folding pattern, suggesting significant structural complexity. This complexity could potentially influence recombination efficiency. Fig 4b compares the monoclonal colony formation under three experimental conditions: CK1 (non-treated), CK2 (heat-shock only, without chilling), and heat-shock treatment (with chilling). Notably, CK1 failed to yield any monoclonal colonies, while CK2 showed limited colony formation. In contrast, the heat-shock treatment, which included a chilling step, led to a substantial increase in monoclonal colony formation, regardless of the vector construction method. These results underscore the critical role of heat-shock treatment with chilling in improving the transformation efficiency of complex recombinant vectors, highlighting its necessity for overcoming the challenges posed by structurally complex DNA.

**Figure 4.**
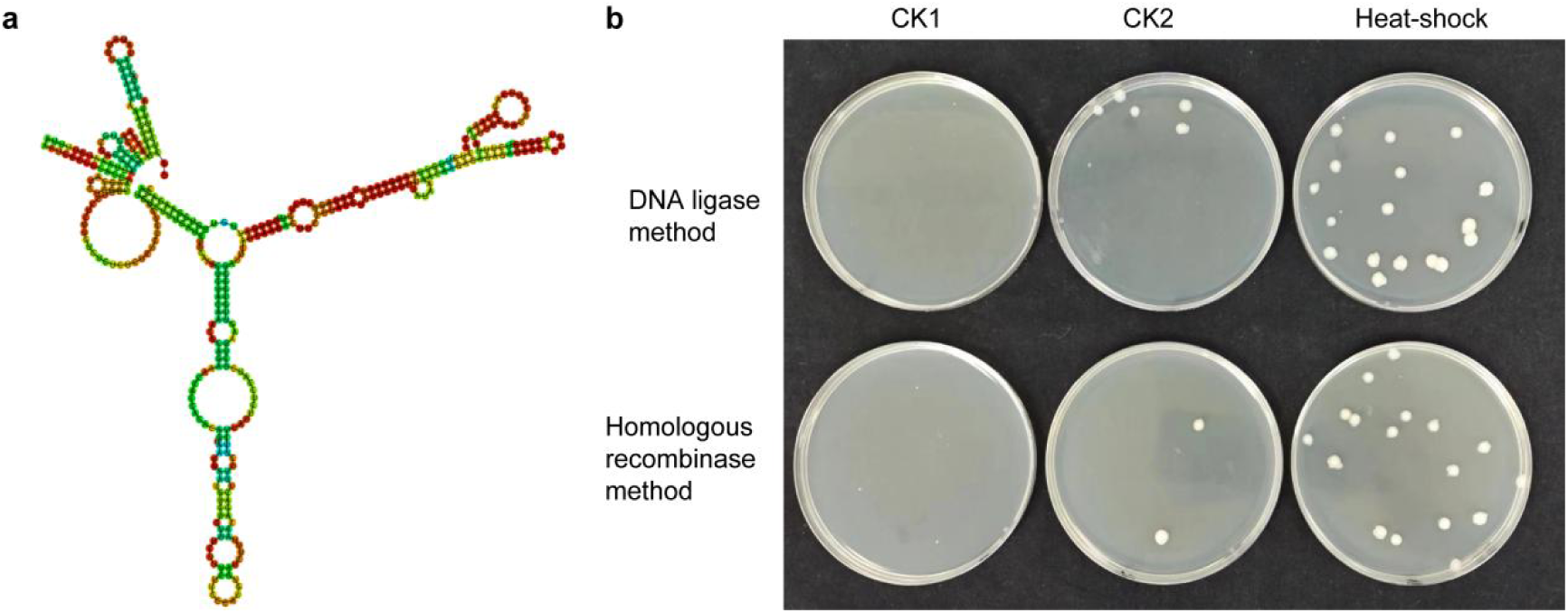
Effect of heat-shock treatment on monoclonal colony formation for complex recombinant vectors (pSYB06). This figure illustrates the impact of heat-shock treatment on monoclonal colony formation for complex recombinant vectors constructed using either the DNA ligase or homologous recombinase method. (a) *EIL1* fragment is predicted with the complex secondary structure using RNAfold. The intricate folding patterns indicate a high degree of structural complexity, which may impact its recombination efficiency. (b) Three conditions were tested: CK1 (non-treated), CK2 (heat-shock only, without chilling), and heat-shock treatment (including chilling). CK1 failed to produce any monoclonal colonies, while CK2 resulted in limited monoclonal colony formation. In contrast, heat-shock treatment with chilling significantly increased monoclonal colony formation for both vector construction methods. These results highlight the critical role of heat-shock treatment with chilling in the transformation efficiency of complex recombinant vectors.

## Discussion

This study provides significant insights into the effects of plasmid conformation and heat treatments on key molecular biology processes, offering practical advancements for DNA manipulation and vector construction. The observation that supercoiled plasmids exhibit higher mobility during agarose gel electrophoresis aligns with previous findings that the compact and twisted conformation of supercoiled DNA enables it to traverse the gel matrix more efficiently than relaxed or linear forms (Sambrook & Russell, 2001). This result highlights the importance of considering plasmid conformation in electrophoretic analysis and experimental design. The enhanced mobility of supercoiled plasmids not only aids in their separation but may also have downstream implications for applications like cloning and transfection. Previous studies have suggested that the compact nature of supercoiled DNA enhances cellular uptake and expression efficiency, making it a preferred choice for these applications (Hirota et al., 2000; Hanahan, 1983).

The increased digestion efficiency observed with heat-treated PCR products further emphasizes the importance of optimizing reaction conditions for enzymatic processes. Heat treatment likely facilitates DNA unfolding or conformational changes, granting better access to restriction enzyme binding sites, as reported by several studies (Smith et al., 1995; Xu & Davis, 2000). This improved specificity and efficiency are crucial for achieving reproducible results in molecular cloning and similar DNA manipulation techniques. Moreover, the enhanced frequency of monoclonal colony formation following heat treatment underscores the role of complete and accurate digestion in improving transformation efficiency.

The heat-shock treatment, particularly when paired with a chilling step, significantly enhances the formation of monoclonal colonies for recombinant vectors, as demonstrated across different vector construction methods. This is especially relevant for complex recombinant vectors, such as those containing the *EIL1* fragment, where structural intricacy often poses challenges to transformation efficiency. Consistent with earlier findings, heat-shock treatment appears to improve DNA fragment alignment and integration by modifying cellular membrane permeability and stabilizing DNA complexes (Dower et al., 1988; Inoue et al., 1990). These findings suggest that this treatment is a universally applicable approach to increasing recombinational efficiency, even in challenging scenarios involving structurally complex DNA.

The strategy of combining heat treatment with immediate cooling represents an efficient and versatile approach for enhancing the success rates of molecular biology workflows. By reducing the structural complexity of DNA and improving the efficiency of PCR product-vector ligation, this method simplifies the construction of recombinant vectors. It holds particular promise for advancing gene cloning, genetic recombination, and gene therapy development. For example, in gene cloning, this treatment could lead to higher yields of correctly cloned sequences, reducing verification and analysis time (Molecular Cloning: A Laboratory Manual, 3rd edition). In synthetic biology and gene editing, the improved efficiency of DNA manipulation could enable the construction of more complex and precise genetic constructs, further advancing the field (Jinek et al., 2012).

Additionally, this technique offers significant potential for the preparation of gene therapy vectors. Efficient vector construction and the stability of therapeutic DNA are critical for ensuring the efficacy and safety of gene therapies (Kay et al., 2001). By improving the integrity of inserted sequences and the overall success of vector assembly, the heat-shock and chilling approach could contribute to the development of more reliable and effective gene therapy strategies.

In conclusion, the findings of this study represent a meaningful advancement in molecular biology techniques. By addressing common challenges in DNA manipulation and vector construction, this optimized method offers a reliable and efficient solution for researchers. As this approach continues to be explored and refined, it holds the potential to become a standard practice in molecular biology laboratories, enabling innovations in synthetic biology, genetic engineering, and therapeutic development.

## Conflict of interest

The authors do not have any conflicts of interest.

## Acknowledgements

Thanks to all team members involved in this study. This work was supported by grants from Wuhan BioRun Biosciences Co., Ltd. to Shu Yuan, and by the Hubei Provincial Science and Technology Plan Project (Grant No. 2025BBA008).

## Notes

### Competing Interest Statement

The authors have declared no competing interest.

## References

Sambrook, J., & Russell, D. W. (2001). Molecular Cloning: A Laboratory Manual (3rd ed.). Cold Spring Harbor Laboratory Press.

Dower, W. J., Miller, J. F., & Ragsdale, C. W. (1988). High efficiency transformation of E. coli by high-voltage electroporation. Nucleic Acids Research, 16(13), 6127–6145.

Green, M. R., & Sambrook, J. (2012). Molecular Cloning: A Laboratory Manual (4th ed.). Cold Spring Harbor Laboratory Press.

Mullis, K., et al. (1986). Specific synthesis of DNA in vitro via a polymerase-catalyzed chain reaction. Methods in Enzymology, 155, 335–350.

Hanahan, D. (1983). Studies on transformation of Escherichia coli with plasmids. Journal of Molecular Biology, 166(4), 557–580.

Jasny, B. R., et al. (1984). Gene cloning in E. coli using plasmids and bacteriophages. Proceedings of the National Academy of Sciences, 81(14), 4311–4315.

Vojta, A., & Tullius, T. D. (1987). Vector construction and gene cloning: Methods and applications. Gene, 55(1), 1–16.

Gibson, D. G., et al. (2010). Complete chemical synthesis, assembly, and cloning of a Mycoplasma genitalium genome. Science, 319(5867), 1215–1220.

Luo, Z., et al. (2002). A new expression vector for protein production in E. coli. Gene, 297(1), 141–146.

Zhang, Y., et al. (2011). A versatile plasmid system for protein expression in multiple bacterial species. Journal of Biotechnology, 151(1), 76–84.

Rosamond, J. L., & Hileman, R. E. (1989). Plasmid-based systems for gene expression in bacteria and mammalian cells. Biotechnology Advances, 7(4), 705–711.

Cohen, S. N., et al. (1973). Construction of biologically functional bacterial plasmids in vitro. Proceedings of the National Academy of Sciences, 70(11), 3240–3244.

Morrow, J. F., et al. (1983). Plasmid vectors for the cloning of genes in E. coli. Journal of Molecular Biology, 166(3), 765–775.

Hirota, Y., Smeaton, M. B., & Van Houten, B. (2000). DNA conformation and cellular uptake efficiency in transfection experiments. Molecular Therapy, 1(5), 343–350.

Smith, C. L., et al. (1995). Optimizing enzymatic cleavage of DNA: Effects of structure and sequence. Nucleic Acids Research, 23(5), 721–727.

Xu, W., & Davis, R. W. (2000). Impact of heat denaturation on restriction enzyme digestion. BioTechniques, 29(3), 624–630.

Inoue, H., Nojima, H., & Okayama, H. (1990). High efficiency transformation of Escherichia coli with plasmids. Gene, 96(1), 23–28.

Jinek, M., et al. (2012). A programmable dual-RNA-guided DNA endonuclease in adaptive bacterial immunity. Science, 337(6096), 816–821.

Kay, M. A., Glorioso, J. C., & Naldini, L. (2001). Viral vectors for gene therapy: The art of turning infectious agents into vehicles of therapeutics. Nature Medicine, 7(1), 33–40.

